# Physiological constraints limiting the growth of *Drosophila* larvae

**DOI:** 10.1101/2023.12.20.572459

**Authors:** Jürgen Schönborn, Irfan Akhtar, Andrea Droste, Stephanie Wesemann, Mathias Beller

## Abstract

The phenotype of organisms is the result of complex interactions between physiology, developmental conditions, and intra-organismic processes, such as metabolism and genetic adaptation. Organisms pursue various developmental strategies that depend on factors such as available resources. How organisms regulate and control critical developmental principles is not fully understood. In this study, we investigated the dynamic development of *Drosophila melanogaster* larvae through *in vitro* and *in silico* experiments. First, we enhanced the larval metabolic network FlySilico and its parameters using growth and metabolite measurements. Subsequently, we established a dynamic flux balance analysis approach and incorporated spatial physiological information from the larval gut to improve the prediction results. We successfully predicted larval growth on different media and developmental critical processes, such as the emergence of larval metamorphosis. This expands the ability to investigate and understand the process of the larval critical weight and, furthermore, the influence of critical developmental processes of multicellular organisms on metabolism. Thus, we present an approach to understand the development, metabolism, and the role of physiology in the development of multicellular organisms.

**Graphical Abstract:** 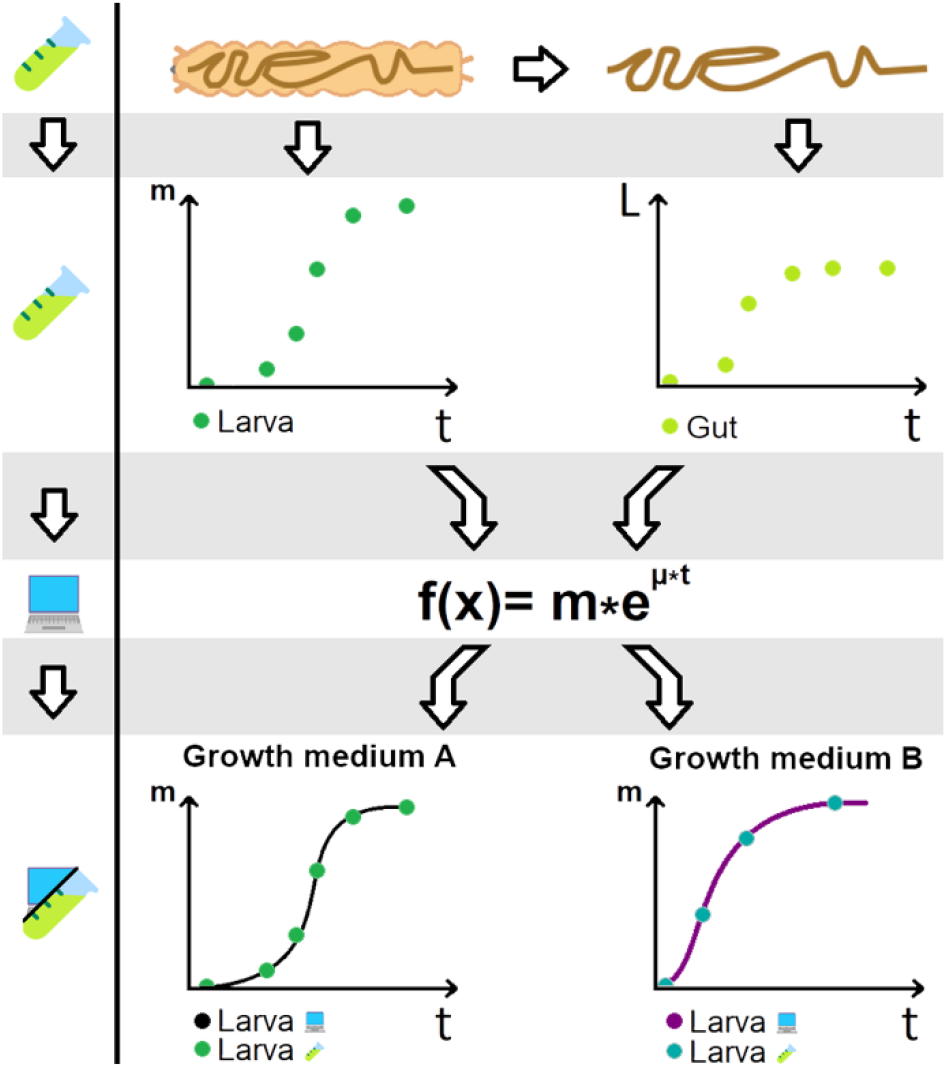

## Introduction

Physiology plays a pivotal role in determining the capabilities and developmental outcomes of all organisms. Influences like environmental conditions affect the physiology of organisms. In the progress of organism’s growth, malnourished higher organisms often manifest smaller overall sizes, leading to reduced organ or tissue proportions when compared to well-fed counterparts (McCance, 1960). The physiology of all living beings is an expression of their phenotypic traits influenced by diverse genetic responses to the varying environmental conditions experienced by the organism (Mitchell-Olds et al., 2007). Even small organisms like *Drosophila melanogaster* exhibit similar behavioral responses to changes in their environment. Throughout the larval growth stage, *Drosophila* undergoes a quick process of cellular division and specialization, during which numerous essential genes involved in development are expressed. The progression of this growth phase can be affected by several factors, such as nutrient availability and temperature, which may alter the timing and pace of developmental events. Given its complexity, the growth of *Drosophila* larvae has been a popular model for comprehending the genetic and molecular mechanisms behind growth and development across various higher organisms (Roberts, 2006; Baker and Thummel, 2007; Melcher et al., 2007). Adult flies that developed from larvae raised at 18 °C show decreased mass and size, but exhibit a higher maximum walking speed compared to those developed at 25 °C on the same growth medium (Crill et al., 1996).

Physiology can be altered through genes and signals, although many of the complex interactions between them are still not fully understood as of today. These interactions, often involving a multitude of biological components and pathways, contribute to the distinctive characteristics and behaviors observed in organisms. While understanding these interactions demands a substantial investment of knowledge of the biological components and resources for investigation (Ginzinger, 2002; Tkačik and Walczak, 2011), the outcome is the emergence of a phenotype uniquely specific to the interactions themselves (Wright, 1941; Shingleton, 2010). These phenotypes are possibly easier to quantify as to measure the underlying gene-signal interaction network that led to the phenotype.

Under conditions of unrestricted feeding and stable environmental conditions, the duration of the larval stages, particularly the third instar larval stage, remain constant. This duration is sufficiently long to accumulate a large reservoir of resources, which is essential for beginning and surviving the metamorphosis (Aguila et al., 2007; Merkey et al., 2011). Larvae need to reach the so called “critical weight” (or “critical size”) (Moed et al., 1999). The critical weight is defined as the minimal weight needed to enter and survive the pupation to become an adult fly (Shingleton et al., 2007). Depending on how nutritious the growth medium is, the larvae can grow larger beyond the critical weight. It is known that growth of larvae and their tissue is dependent on the interaction of signal hormones (Riddiford and Ashburner, 1991). In addition, more known and unknown signals, along with genes, interact in various ways towards a complex gene-signal network that enables the growth in a specific environment. Despite the complexity of the mechanics behind growing in different environments all organisms show distinct phenotypical and morphological features. Those features can be used to derivate the progress of growth to pin down growth phases and their underlying features.

The understanding of the underlying mechanisms of larval growth can be examined in different ways. Wet lab experiments can be conducted to investigate larval growth in terms of size and weight or to explain the genetic factors underlying larval development. The identification of such genetic factors offers valuable insights into the understanding of development but requires extensive labor and complex analysis to obtain meaningful conclusions (Anholt and Mackay, 2004). Furthermore, *in silico* experiments offer a valuable approach to gaining insights into the progression of larval growth during development. For this reason, different approaches can be used. One way to perform meaningful *in silico* experiments is the use of flux balance analysis (hereafter referred to as FBA) in combination with wet-lab experiments. *In silico* experiments with FBA were already successfully used with various organisms: (i) *Escherichia coli* was used to investigate growth, overflow metabolism, and energy consumption (Kauffman et al., 2003; Orth et al., 2010; Zeng and Yang, 2019). (ii) The growth and ethanol production of yeast cultures (Förster et al., 2003; Hjersted and Henson, 2009). (iii) The ATP yield of different carbon sources for human (Swainston et al., 2016). Moreover, FBA was already successfully used to understand the development and environmental impacts on *Drosophila melanogaster* larvae and flies by examining resource allocation, impact of altered growth conditions and the resulting impact on metabolic fluxes (Coquin et al., 2008; Feala et al., 2009; Schönborn et al., 2019; Cesur et al., 2023).

Investigations were made concerning the dynamics of organism’s growth, changes in metabolism, nutrient uptake, and metabolite excretion over time using more advanced techniques involving FBA. Dynamic flux balance analysis (hereafter referred to as “dFBA”) is one extended version of FBA that allows the investigation of dynamic changes in the simulation conditions, including extracellular concentration changes (Hjersted and Henson, 2006; Meadows et al., 2010; Henson and Hanly, 2014). dFBA enables the prediction of changes in metabolite concentrations and fluxes both intra- and extracellularly over time. This was enabled through incorporating constraints that change the flux rates over time. This approach has found extensive use in predicting dynamic growth in cell culture dynamics making it suitable for studying cellular metabolism in batch cultures, for example for *E. coli* (Varma and Palsson, 1994) or Yeast (Hjersted and Henson, 2006).

This method illustrates that predicting the growth dynamics of cell batch cultures is feasible, although not all organisms share the same growth conditions as those found in batch cultures. For instance, laboratory-reared *Drosophila melanogaster* grow in an environment that closely resembles a continuous culture growth, where they receive a constant and steady supply of nutrients throughout their entire development with no changing extracellular metabolite concentrations. Therefore, the dFBA approach must be adjusted to effectively predict and describe the growth of organisms like *Drosophila* larvae. This can be achieved by integrating growth-related attributes to the simulations, such as the physiological properties of organisms.

We previously published the curated metabolic network FlySilico (Schönborn et al., 2019) of *D. melanogaster* based on time-resolved growth and metabolite measurements from larvae grown on *holidic diet* (Piper et al., 2014) (hereafter referred to as “HD”). The metabolic network comprises the metabolic pathways of the core metabolism of *Drosophila*. To comprehensively explore larval metabolism across various environmental conditions a biomass function to simulate larval growth was formulated based on experimental metabolite measurements. However, FlySilico, predominantly characterizes the phase of larval growth marked by exponential expansion and potentially overlooking critical insights at the onset and conclusion of larval development.

Considering these limitations, we present an enhanced version of FlySilico. Our goal was to refine predictive accuracy by incorporating the initial and final stage of larval development, with a focus on the larva-to-pupa transition. Additionally, we aimed to identify factors that impose constraints on growth during these crucial developmental phases. First, we improved and expanded the underlying data that are used to determine growth parameters based on experimental results. Secondly, we constructed a dFBA approach that fits the prediction of larval growth over the time. Thirdly, we show that phenotypes such as morphological attributes can be used to explain the growth through the larval stage of wild-type *Drosophila melanogaster*. We investigated the growth and pupation timing in dependence of their growth environment by using physiological information (such as size and weight) with a dFBA approach. For that we examined the larval gastrointestinal tract (hereafter referred to as “gut”) and the larva itself on chemically defined and undefined growth media. Our data provide confirmation that the interplay between physiological characteristics and metabolic processes can effectively be used to estimate both the timing of pupation in larvae and their metabolic patterns during growth.

## Results

### Detailed analysis of larval growth and development

Our decision to acquire a more extensive dataset was motivated by the observation that the previously published FlySilico dataset lacked coverage of both the initial and final stage of the larval development. It had the potential to overlook significant developmental phases, such as the critical pupation timing. The pupation timing marks the completion of larval development. The inclusion of data about crucial developmental phases assures to improve the precision and quality of insights into larval development.

Under typical conditions, characterized by sufficient nutrient availability, stable temperature, and a wild-type genetic background, the weight increase of developing *Drosophila melanogaster* larvae follows an S-shaped exponential growth pattern, commonly referred to as sigmoidal growth. This growth pattern is evident in our dataset (Fig. 1). The data reveal a proportional relationship between the larval growth and the metabolite content, presenting three distinct phases. Initially, the growth progresses moderately, resulting in a lag-like phase. Subsequently, an exponential growth phase occurs, which leads to a third phase where the growth reaches a plateau which ultimately leads to pupation. We measured different larval growth parameters (e. g. larval weight and larval size dimensions) and the metabolic composition in a much more fine-grained and time-resolved manner in comparison to the first data set that we used in the first published FlySilico version (Schönborn et al., 2019).

**Figure 1:**
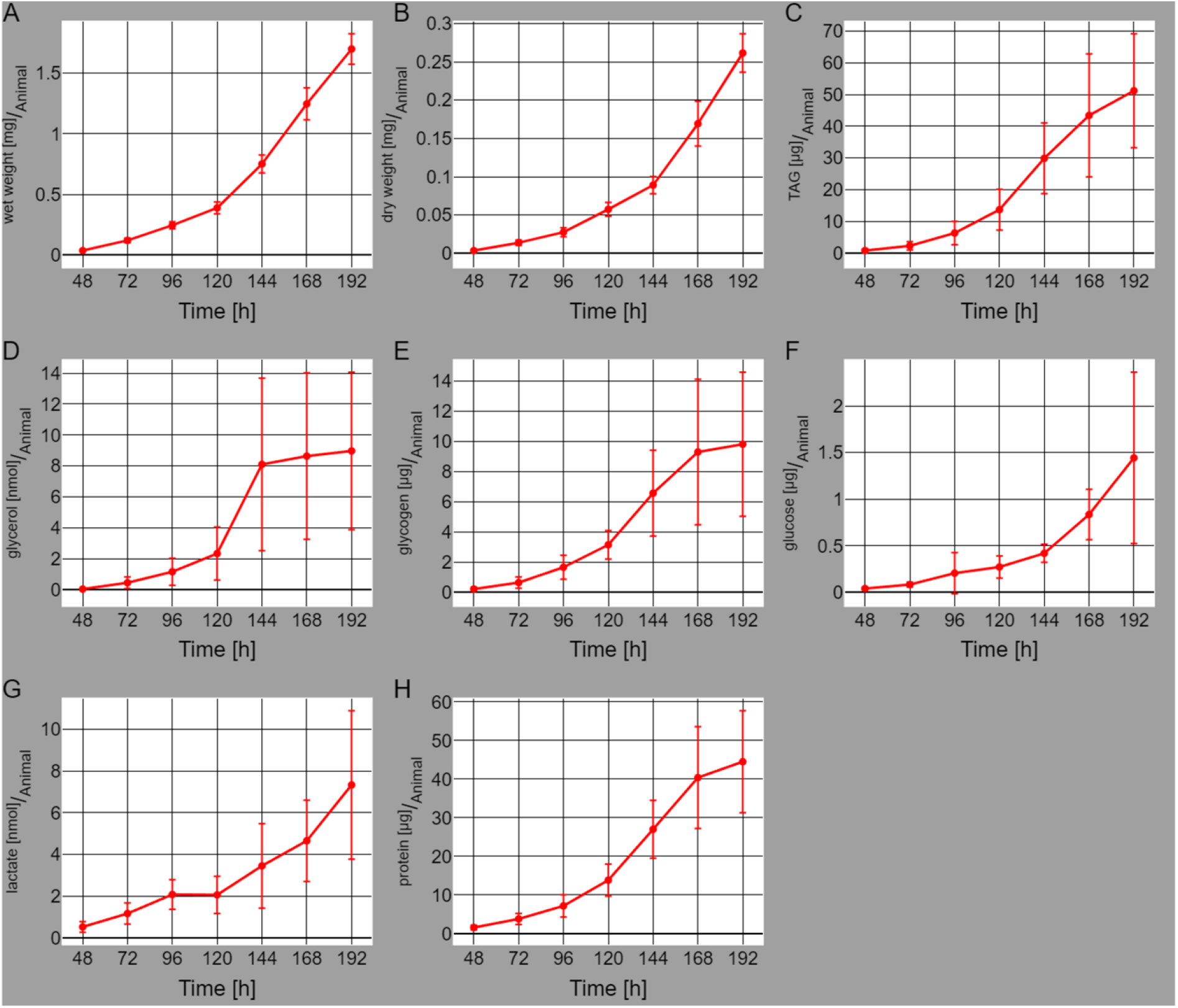
Metabolic profiling of the larval developmental of *Drosophila melanogaster* with growth from 48 to 192 hours after egg laying on holidic diet (HD). Wet weight (A) and dry weight (B) measurements of larvae from 48 hours every 24 hours to 192 hours. (C–H) Absolute quantification of TAG, glycerol, glycogen, glucose, lactate, and triglyceride (TAG) levels. The data is presented as mean values ± standard deviation normalized to the number of animals per sample (triplicate measurements). On average the pupation started around the 192 hours mark.

### In-depth enhanced larval *Drosophila* metabolic model

We expanded the previously published metabolic model FlySilico (Schönborn et al., 2019) by incorporating different pathways in greater detail such as the fatty acid biosynthesis and purine and pyrimidine metabolic pathways (Supplementary Table 1). These additional reactions cover the synthesis of macromolecules and polypeptides used to build up important tissues and structures. The current version of FlySilico contains 584 reactions and 326 unique metabolites (Table 1).

**Table 1:** Comparison of the previously published FlySilico and the FlySilico version from this study. The number of compartments lists the number of cellular compartments present in the respective metabolic network. The count of biomass components denotes the individual metabolite components form the biomass equation of the respective metabolic network.

| Model | FlySilico 1.0 | FlySilico 2.0 | $\Delta$ Increase |
| --- | --- | --- | --- |
| # Reactions | 363 | 583 | +220 |
| # Metabolites | 203 | 326 | +123 |
| # Genes | 261 | 443 | +182 |
| # Compartments | 3 | 6 | +3 |
| # Biomass Components | 31 | 40 | +9 |

The more detailed fatty acid biosynthesis pathway representation was designed to allow us to make more precise statements on how the fatty acids are utilized in the metabolism and in terms of their impact on larval growth, as the fat body of larvae is one of the most important organs for growth and survival for the organism (Zheng et al., 2016). The fat body enables to store as much fatty acids as needed to survive the larval metamorphosis.

*Drosophila* larvae show a sigmoid weight gain over the course of the development (Fig. 1A, B). This comes with the problem that the growth rate and therefore the biomass function changes over the course of developmental timing. To address this issue while maintaining the use of flux balance analysis to predict the growth we chose to calculate a weighted mean metabolite coefficient for defining the demand of each metabolite essential for growth (view method section). This allows us to use a single coefficient for each metabolite contained in the biomass function over the time of development. Besides the improved single coefficient for each metabolite in the biomass function we incorporate different important macromolecules into the biomass formulation.

Important structures for larval survival are the mouth-hooks and cuticle which both are made of a protein-chitin-matrix. The mouth-hook functions as the tool to take up nutrients into the organism while the cuticula functions as a stabilizer and protection for the larval body (Chihara et al., 1982; Ashburner, 1989). We include the metabolite chitin into the biomass function based on data from literature. The chitin content of an adult fly is around 7.85 % (Kaya et al., 2016) which we used as an approximation for the content of chitin in a larva.

RNA and DNA play an important role in the growth of every organism by applying significant influence on various aspects of an organism’s development (Church and Robertson, 1966; Chícharo and Chícharo, 2008). For instance, anomaly in nucleic acids can contribute to the growth of tumors or other genetic mutations (Loeb et al., 2003). Since the previously published version of FlySilico does not contain any sufficient representation of nucleic acids, this absence limits the ability to make more precise observations. Therefore, meaningful conclusions about the impact and alterations of genetic information during development cannot be made without implementation. Thus, the inclusion of nucleic acids in our modeling is of importance to enhance the precision of the simulation results. We included nucleic acids to the metabolic network in a simplified form. The underlying building blocks of nucleic acids are incorporated into the biomass function. The building blocks are dAMP, dCMP, dGMP and dTMP for DNA and AMP, CMP, GMP and UMP for RNA. All mentioned building blocks received a biomass coefficient based on literature values from the yeast model of Förster et al. (2003). This should sufficiently enhance the predictive power of the larval metabolic network.

### Modeling dynamic *Drosophila* larval growth

Flux balance analysis is usually used to predict a flux distribution that minimizes or maximizes a defined objective function (Varma and Palsson, 1994). This objective function can be a biomass function which describes the needed metabolites to synthesis biomass of an organism or a flux that synthesis a metabolite of interest (Kauffman et al., 2003). These predictions are performed by FBA for a steady-state system. To predict the whole development of an organism like a *Drosophila* larva, the FBA approach needs to be calculated through developmental time which represents a non-typical steady-state system. This can be achieved using dynamic flux balance analysis (dFBA) modeling, where the prediction is performed in a time dependent manner with changing conditions. This technique has been previously demonstrated to successfully predict the growth of *E. coli* within a batch culture (Varma and Palsson, 1994). In a batch culture, *E. coli* relies on initial metabolite concentrations, which gradually evolve over time until all growth-supporting metabolites are consumed.

*Drosophila* larvae usually do not grow in a batch culture. The larval growth in a laboratory culture occurs usually under constant/overflow metabolite concentrations. The growth of larvae is different to a batch culture growth and has three important key setups: Firstly, the growth medium is usually a homogenous mixed system. Secondly, the metabolites used to grow do not deplete through the development from larva to an adult fly. Thirdly, no significant amounts of metabolic end products accumulate. This growth resembles rather the attributes of a continuous growth culture. The continuous culture growth is described as growth in an environment with steady stream of nutrients which allow growth at a continuous rate over an indefinite time (Novick and Szilard, 1950).

As a result, dFBA that models the growth and changing metabolite concentrations in larval metabolism throughout development, indicates unrestricted growth for a *Drosophila* larva (Supplementary Fig. 1).

Our hypothesis was that a fundamental mechanism limits the unrestricted growth of wild-type *Drosophila* larvae and contributes to the checkpoint regulating the entry into pupation. Therefore, we conducted an analysis of larvae and their various tissues, as the physiological characteristics could manifest the fundamental mechanism in a phenotypic manner (Fig. 2). Upon analyzing the larval gut, we observed that the spatial parameters governing gut growth on HD growth medium halt before the larva transition into the wandering L3 stage, marking the onset of pupation (Fig. 3). The termination of gut growth prior to the wandering phase and pupation, presents an opportunity to establish a constraining parameter. This parameter endorses the simulation of larval growth to mimic the natural development observed in the real world.

**Figure 2:**
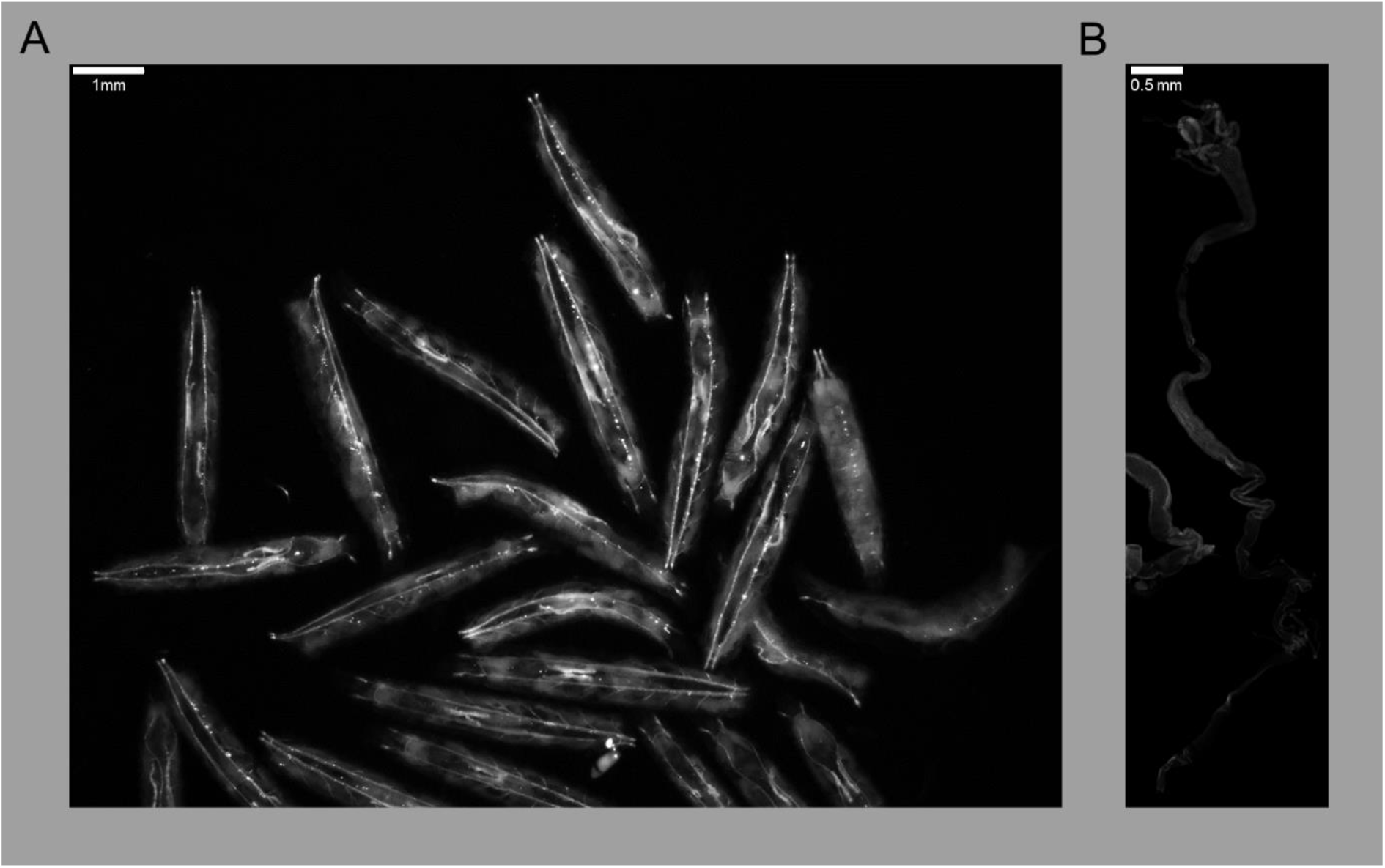
Growth of larvae reared on chemically defined medium. (A) Larvae growing on HD medium at 168 hours AEL. The photo was taken with a Stereomicroscope from Zeiss (SteREO Discovery.V8). The length, width and area were set manually by using the software Zeiss Zen 2.3 lite (blue edition). (B) Example of a dissected larval gut from an individual raised on HD at the 192 hours timepoint AEL. Sample was imaged with an Operetta CLS high content screening microscope (Perkin Elmer) and were recorded with a 20x air objective.

**Figure 3:**
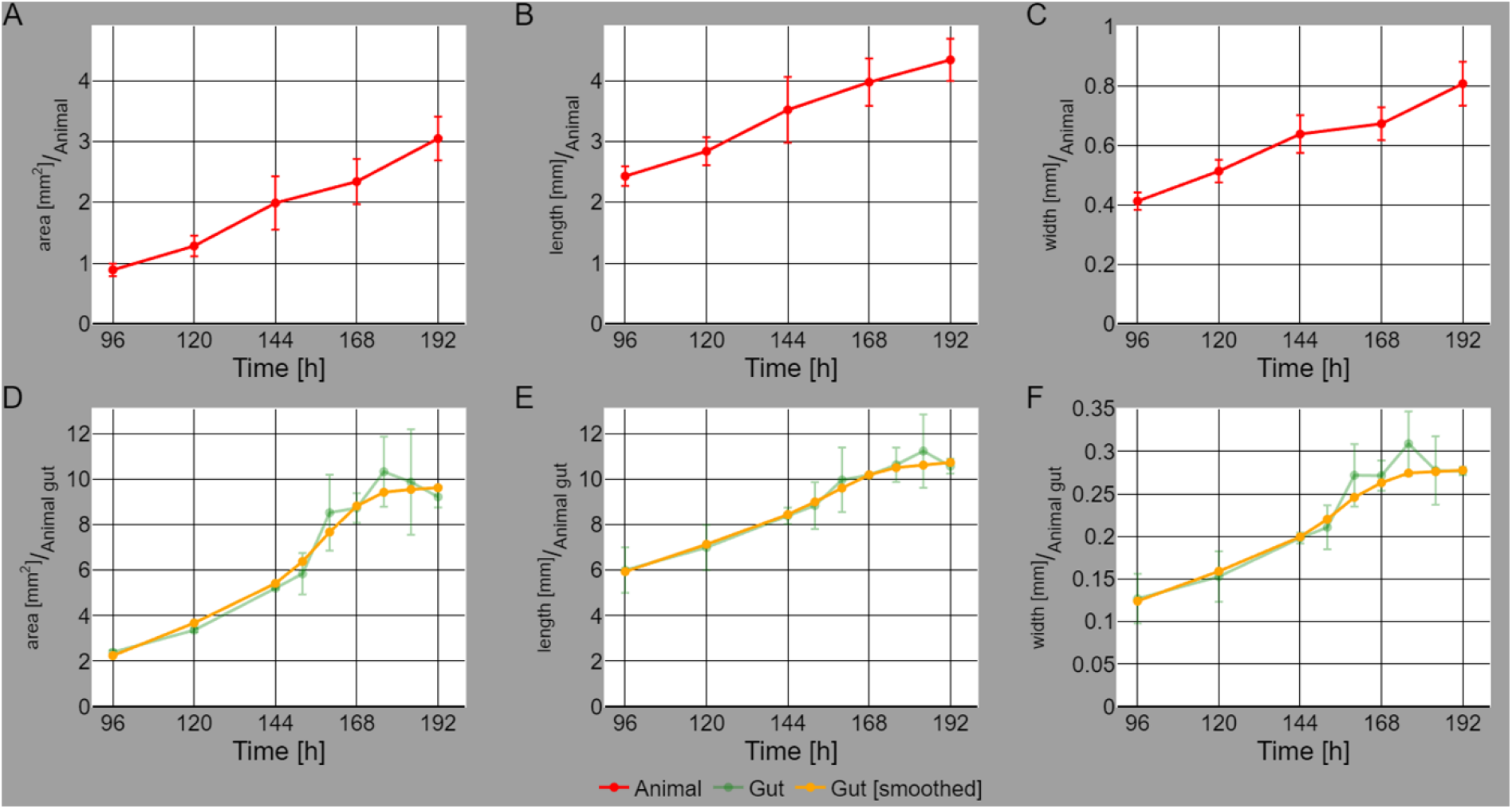
Growth dynamics of *Drosophila melanogaster* larvae and gut. (A-C) Gradual growth in the area, length, and width of developing larvae. The area is calculated based on the length and width of a larvae. Each timepoint consist of at least 20 larvae (triplicate measurements). The data are presented as mean values ± standard deviation normalized to the number of animals per sample. (D-F) Increase of area, length, and width of the gut from developing larvae. The area is calculated based on the length and width for each gut. Each timepoint consist of at least 8 guts (triplicate measurements). The unsmoothed data (shown in green) and smoothed data (shown in yellow) illustrate the sigmoid progression of larval gut growth. Initially, a rapid growth from 96 hours to 168 hours is present which is followed by a nearly plateau from 168 to 192 hours. The data is presented as mean values ± standard deviation normalized to the number of guts per sample (green). On average the pupation started around the 192 hours mark.

### Modeling the gut as a constraining parameter in growth simulations

To address the previously mentioned issue of unlimited growth in larvae, we implemented a penalty function based on morphological and physiological attributes of the larval gut. The design of this penalty function was inspired by the sigmoidal growth patterns observed in both larval weight (Fig. 1A and B) and gut development (Fig. 2A-C).

The penalty function was implemented to regulate nutrient uptake through the gut and subsequently limit larval growth (see Method section). During the early stages of larval development, the penalty function did not impose any limitations on growth, as the gut efficiently absorb nutrients from the growth medium and allocate the resources fully to larval growth (Fig. 4; Supplementary Fig. 3). As the development further progresses a change in the growth pace is visible which is the result in a decrease of growth rate due to the limitation of the gut function on nutrient uptake (Fig. 4).

**Figure 4:**
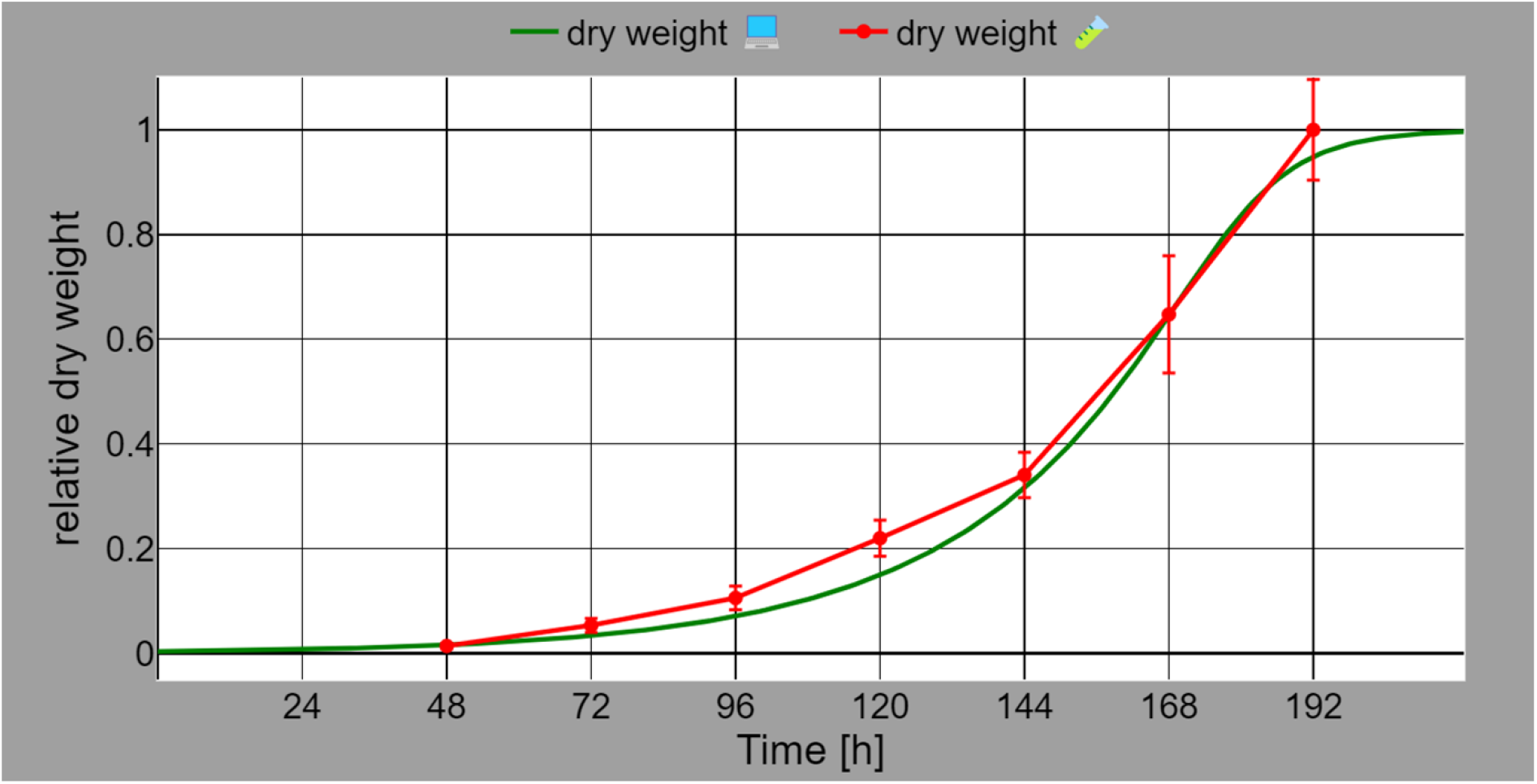
Comparison of *in vitro* and *in silico* larval growth on chemically defined growth medium. We simulated the *in silico* growth of a larva (green; dry weight) on HD by using the dFBA approach (see method section). The growth of the larva resembles a sigmoidal growth curve till it reaches a plateau starting around 192 hours. The growth of larvae reared *in vitro* (red; dry weight) on HD, starts at 48 hours till 192 hours AEL. The growth *in vitro* shows a small variance in comparison to the *in silico* growth based on the relative dry weight values. On average the pupation started around the 192 hours mark.

This transition is evident during the phase when larvae enter the critical weight range, leading to a stage of minimal to no growth. Simultaneously, growth of the gut declines and its expansion halts (Fig 3). The penalty function mimics this process as it reduces the nutrient uptake specifically for growth in the simulation. This reduction in nutrient uptake resulted in a decrease in growth rate. As the growth rate continued to decrease, it reaches a point where no further growth was possible. Upon reaching this point, the critical weight is already achieved and, along with the halting of gut growth, it serves as a signal of the larva’s transition towards eventual pupation.

The penalty function is mathematically described as a negative sigmoidal function (Supplementary Fig. 3) which runs between 1 and 0. This function is based on different attributes of the larval development which enables the successful integration into the dFBA simulation. The penalty function describes the timing where the larval gut from larvae grown on HD stopped growing. This is roughly around 180 hours after egg laying (AEL) on HD with a function value of 0.5. This is represented by the largest change in the gradient of the function on the timepoint 180 hours AEL which is the turning point of the function (Supplementary Fig. 3). This important time point in the development of larvae under standard conditions grown on HD show a correlation to the critical weight and the end of the development of the gut. Around the turning point of the function a phase is visible which overlaps with the range where the critical weight is reached. This phase of the function starts around 128 hours with a function value of around 1 and ends around 218 hours with a function value of nearly 0. The most crucial impact on the simulation becomes apparent when we observe a particular gradient value, approximately 0.6. This occurs during the time period between 168 hours and nearly 192 hours (Supplementary Fig. 3). Notably, this specific time interval corresponds to the critical weight range, a phase where larvae are capable of initiating pupation. This gradient change results in a sharp decline of the gut function and consequently flattening of the larval growth curve (Fig. 4). This flattening signifies a critical transition point in the larval development process due to the reduced nutrient absorption of the gut.

In summary, the introduction of a negative sigmoidal penalty function based on physiological attributes effectively limits nutrient uptake and controls the growth rate in larvae. This regulation played a crucial role in determining the larva’s entry into the wandering L3 phase and subsequent pupation based on the simulation results (Fig. 4). Having successfully run the dFBA simulation of larval growth on HD under standard conditions by incorporating the penalty function (Fig. 4), it is now intriguing to explore whether the current dFBA simulation can accurately predict outcomes in unfamiliar scenarios.

### Dynamic growth model verification

The first step in our model verification procedure was to select an unfamiliar scenario for the simulation. We chose to test our simulations on a complex medium called low sugar diet as this diet prominently alters the developmental timing (see method section; hereafter referred to as “LSD”). We defined the underlying growth medium parameters by defining the mediums components and their corresponding amounts (Supplementary Table 2). As LSD is a complex medium, the amount and composition of nutrients available for intake into the metabolic network can vary depending on the complex medium defined for the simulation. This variation in the case of growth, can lead to differences of growth rate in the simulation. Next, we performed growth simulations that were promising as the growth timing was predicted to be much shorter (120 hours for larval grown on LSD in comparison to 192 hours for larval grown on HD).

In the case of the complex LSD medium, the penalty function revealed a shift in the turning point to approximately 100 hours AEL (Supplementary Fig. 4). This is due to the accelerated growth rate in comparison to HD medium. The time point of 100 hours shows a function value of 0.5. The gradient is changing in a range which starts around 76 hours till the end of around 128 hours. This is a smaller phase where the gradient changes in comparison to the penalty function for the HD and results from the much faster growth of the larva on LSD.

Subsequently, we performed corresponding *in vitro* growth experiments following the initial simulations carried out. We grew larvae on LSD under the same standard conditions as they were grown on HD (see Method section). We measured necessary parameters like weight for larvae reared on LSD (Fig. 5). We compared our *in silico* simulations to the experimentally assessed data and found both were in accordance with each other proving the robustness of our *in silico* simulations (Fig. 6). The growth simulation indicated that larvae raised on the LSD medium reveal a developmental period of around 120 hours AEL, mirroring the *in vitro* findings, which also report a 120-hour developmental duration for the larvae.

**Figure 5:**
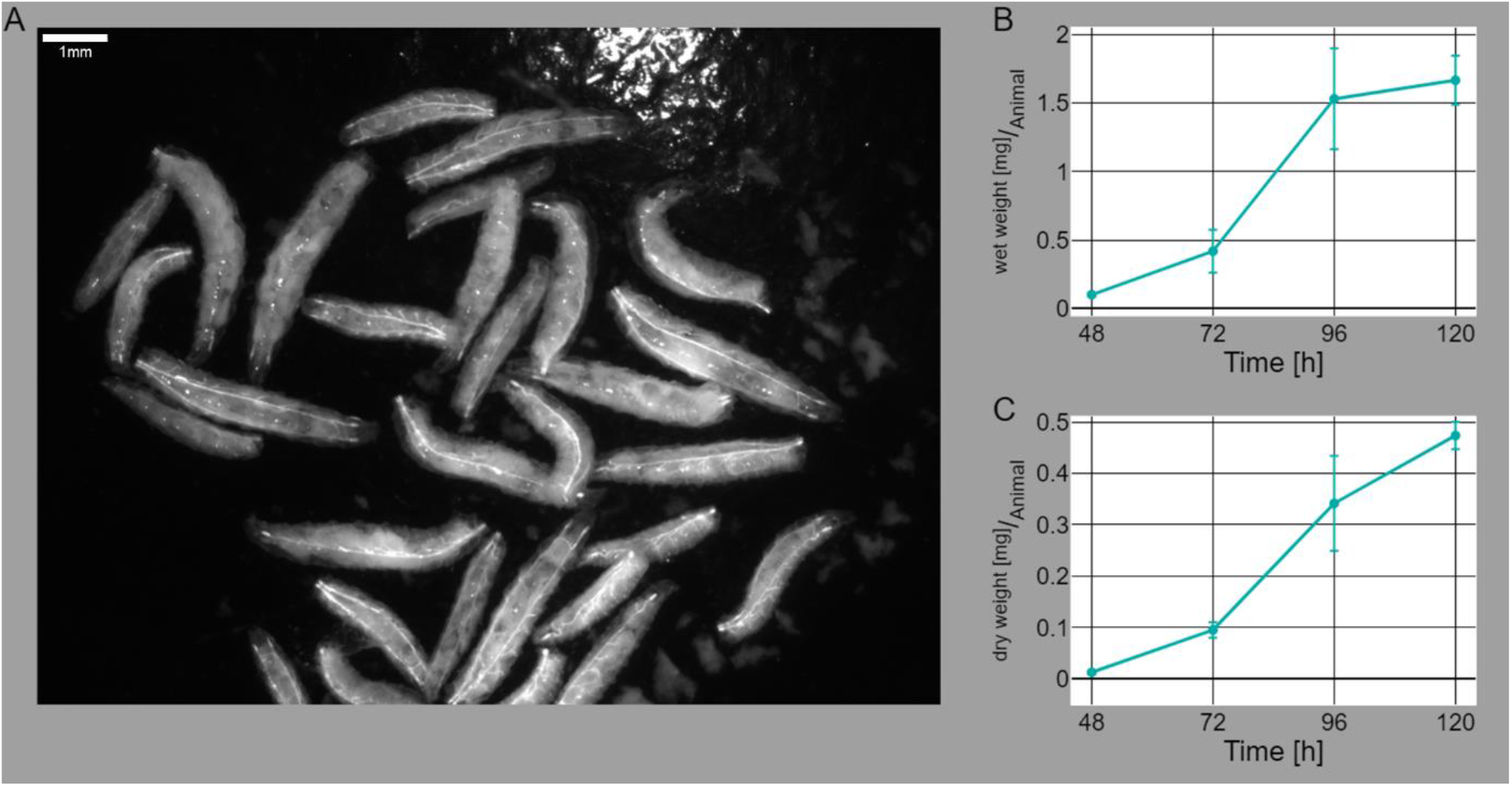
Growth of larvae reared on complex medium. (A) Growing larvae on LSD medium at the timepoint of 72 hours AEL. The photo was taken with a Stereomicroscope from Zeiss (SteREO Discovery.V8). The length, width and area were set manually by using the software Zeiss Zen 2.3 lite (blue edition). Wet weight (B) and dry weight (C) measurements of larvae from 48 hours every 24 hours to 120 hours grown on LSD. Each timepoint consist of at least 50 larvae (duplicate measurements) up to 75 larvae (triplicate measurements). On average the pupation started around the 120 hours mark.

**Figure 6:**
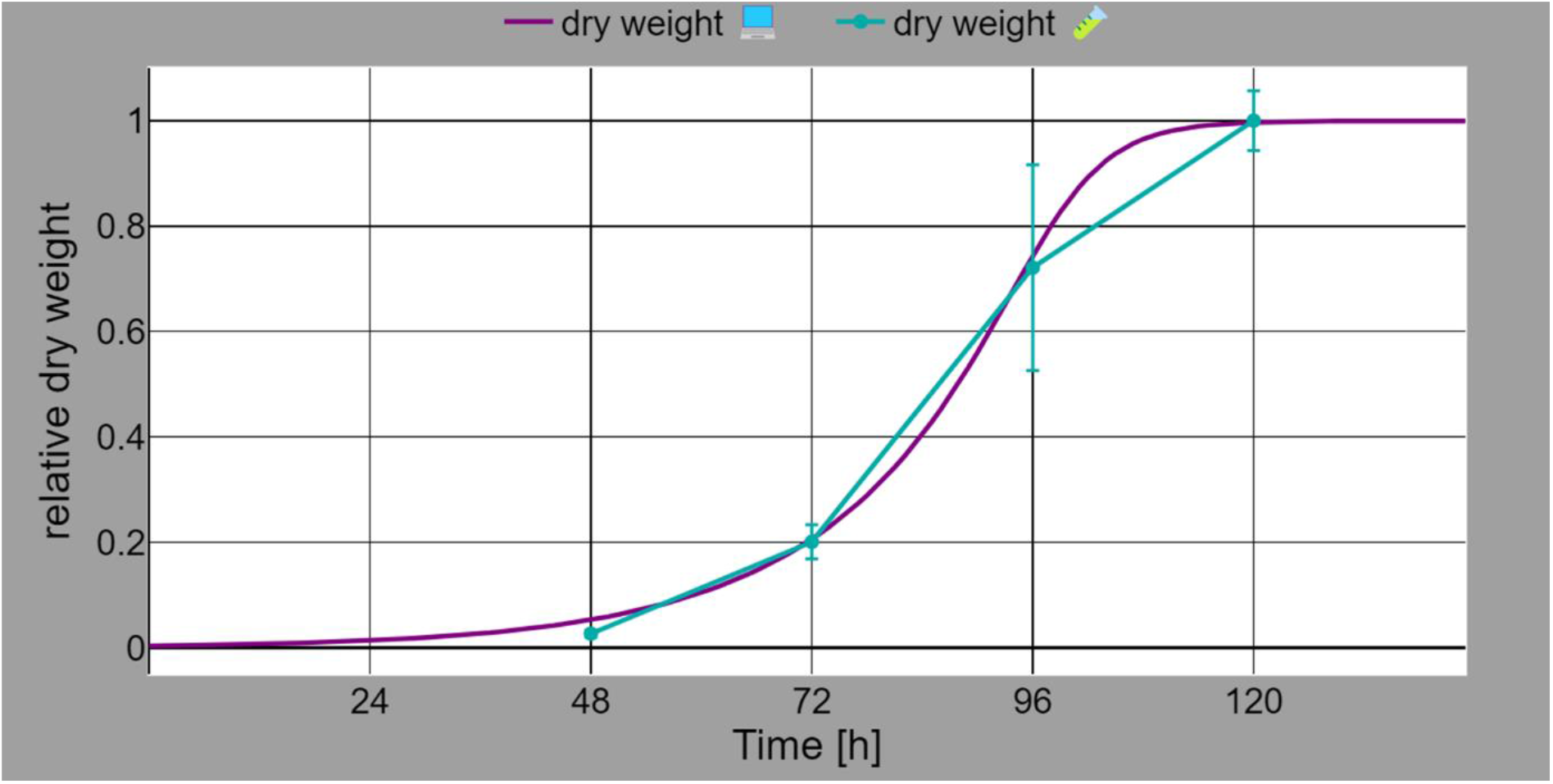
Comparison of *in vitro* and *in silico* larval growth on complex growth medium. We simulated the *in silico* growth of a larva (purple; dry weight) on HD by using the dFBA approach (see method section). The growth of the larva follows a sigmoidal growth curve till it reaches a plateau starting around 120 hours. Larval growth observed *in vitro* (dark cyan; dry weight) on HD medium, starts from 48 hours till 120 hours AEL. The growth *in vitro* shows a small variance in comparison to the *in silico* growth based on the relative dry weight values.

These results demonstrate the consistency between our modelling data and experimental observations, affirming the validity of employing physiological data in combination with dFBA to predict larval development, even in complex environmental conditions.

## Discussion

In this study, we investigated the physiological aspects of *Drosophila melanogaster* larval growth through a combination of *in vitro* experiments and *in silico* simulations. Laboratory-reared wild-type Oregon-R larvae were raised on chemically defined and complex growth medium. Subsequently, we collected physiological and metabolic data, which served as the basis for our *in silico* growth and metabolism modelling analyses.

We translated a fine-grained metabolic profile of growing larvae on HD (Fig. 1) into single coefficients for each metabolite in the biomass function. The growth behavior of *Drosophila* larvae is evidently complex, and a simplistic approach of employing single coefficients for each metabolite in the biomass function falls short of providing a comprehensive explanation for their growth and metamorphosis dynamics. Nevertheless, streamlined models such as FlySilico, with simplified parameters, offer the advantage to understand fundamental principles and mechanisms of metabolic processes and their consequences. These models are useful for identifying larger metabolic pathways to answer specific biological questions. In the context of larval growth, FlySilico demonstrated which metabolic pathways are essentially required for simulating growth (Schönborn et al., 2019). These models can assist in identifying critical factors and interactions within an organism’s metabolism, such as the influence of different nutrients and their quantities on metabolism and growth, as demonstrated in FlySilico (Schönborn et al., 2019). Additionally, in cases where information and data are limited, such models can still provide meaningful insights, whereas large-scale complex models might require additional data. The development of a *Drosophila* larva has different distinct stages (Bate and Martinez Arias, 1993). The developmental process begins with the embryonic stage and progresses through the first, second, and third instar larval stages. Each of these stages show distinct growth patterns and timelines. At the end of the larval development, the larva undergoes a pupation and emerges as an adult fly. The various stages of larval development are expected to have diverse metabolic patterns, much like what is observed during larval embryogenesis (An et al., 2014). The variability in larval growth within a population is noticeable, with some individuals show lower or higher growth rates than others. This variance can be attributed to various factors, such as environmental conditions or genetic backgrounds. To ensure the robustness of conclusions the results are based on the average growth pattern observed in the larval population. Therefore, as distinct metabolic profiles and growth trajectories are expected during different developmental stages, it remains feasible to employ singular coefficients within the biomass function. This approach yields an averaged growth rate over the entire developmental period as these result from the utilization of weighted-average metabolic coefficients. This simplified modelling approach enables the investigation and analysis of larval growth throughout the entire development in simulations with minimal variation to the real world (Fig. 4 and Fig. 1).

Organ growth is influenced by various factors, such as environmental conditions, different signaling pathways, and communication between organs (Andersen et al., 2013). For that reason, an addition of the FBA simulations is needed to simulate these complex signaling pathways. The incorporation of signaling pathways within FBA models help to understand how signals affect metabolic fluxes, as signaling pathways often regulate metabolic processes (Ward and Thompson, 2012). Incorporating these results could significantly impact the accuracy of FBA simulations, leading to predictions that better mirror real-world scenarios in the best case. Although there are established methods that combining (d)FBA with signaling networks (Papin and Palsson, 2004; Price et al., 2004; Covert et al., 2008), these approaches often face challenges due to the requirement for extensive data. These data are not always readily available or sufficiently detailed for use with FBA (Covert et al., 2001; McKechnie et al., 2010; Hiruma and Kaneko, 2013; Nijhout et al., 2014). The approach we used in this study therefore does not delve into complex signaling pathways. To address these challenges, we have chosen a simpler approach: we assume that the signals and interactions governing organ growth led to specific, defined phenotypes. In other investigations, it has been demonstrated that the physiological functions of various organs contribute to the attainment of developmental stages. For instance, Mirth et al. (2005) observed that the larval organ, the prothoracic gland, plays an important role in determining the critical weight required for the metamorphosis of *D. melanogaster* and secretes growth-related hormones. These characteristics can be effectively translated and integrated into our dFBA approach, making it a more practical and accessible method. It is known that (d)FBA is sensitive to the chosen parameters to a certain degree (Varma and Palsson, 1995). Considering these observations, it becomes evident that the introduction of additional parameters, such as a complex signaling pathway, could significantly alter the predictive outcomes. The prediction of larval growth becomes challenging under such conditions.

The *in silico* approach FBA was improved further into a variation of dFBA. The dFBA approach allows simulations were the larval growth and their metabolic content itself can be analyzed during larval development. The dFBA approach is already established and successfully used for different organisms, such as *E. coli* (Varma and Palsson, 1994). It is an established algorithm that is also implemented in different tools such like the COBRA Toolbox (Heirendt et al., 2019). The reason why we created an altered version of dFBA is, that the growth of larvae is not possible to be predicted by the unaltered algorithm. The growth of larvae has fundamentally different conditions to a growth scenario for a bacterial culture. The larval growth medium is homogenous mixed and the used metabolites from the medium usually do not deplete through the larval development. Therefore, the metabolite concentration in the medium does not change which is the fundamental condition that the established dFBA algorithms uses as the metabolite concentration changes over the developmental time are calculated (Varma and Palsson, 1994). This results in the case of the larval development into an unlimited growth (Supplementary Fig. 1) which needed to be addressed. To address this issue for dFBA we analyzed different tissues as we hypothesized that physiological properties can be found to solve this unlimited growth issue. We found that the larval gut is a promising candidate to solve the unlimited larval growth.

The larval gut plays a major role in systemic growth. If larvae sense conditions of poor nutrient concentration of the growth medium, the gut crosstalks through signals with the fat body which signals the reduction of larval growth rate (Shin et al., 2011; Storelli et al., 2011; Andersen et al., 2013). Conversely, this signaling activity is not fully understood. In developing larvae, we observed that the larval gut stops to increase in size earlier than the onset of pupation (Fig. 3). As the larval gut plays an important role in larval growth, it provides an entry point for integration into the modelling of the larval growth. The growth of the larval gut and its premature stop were consequently translated into a growth-determining parameter (see Result section) within the *in silico* simulation. This made the prediction of larval growth over the time, including its termination and the correct pupation timing prediction possible (Fig. 4).

Our early results regarding the predictions of *Drosophila* larvae growth on HD through additional physiological and metabolic data are promising. They suggest that simulations with the enhanced model should be able to correctly predict different conditional scenarios.

To ensure the reliability of the growth-determining parameter, additional *in silico* simulations were performed. In these simulations, the growth for larvae raised on the complex medium LSD were predicted. We reasoned that the successful prediction of the HD growth situation should also elevate the simulations to be able to predict different growth situations of experimentally different conditions. If this is the case, this shows that our approach, to use physiological information for prediction, is valid to be used in changing growth conditions and successful predict those. Based on the previous *in silico* simulations we performed equivalent *in vitro* wet lab experiments on LSD medium (Fig. 5). The simulations revealed that the growth of larvae could be accurately predicted as they align well with the experimental data (Fig. 6). This confirms our assumption that the combination of physiological and metabolic attributes can be used to successfully predict the growth of *Drosophila* larvae on defined and complex growth media.

### Shortcomings of predictions based on phenotypical attributes

The presented results show promising findings to understand the larval growth. The performed experiments and simulations show valid results for wild-type OregonR larvae. It is evident that the case of non wild-type larvae is not precisely predictable as the metabolism of the non wild-type can heavily change in comparison to the metabolism of wild-type larvae. Predicting larval growth from mutation-harboring *Drosophila* lines is especially challenging, as these mutations can induce growth-affecting phenotypes (Migeon et al., 1999). Further iterations and improvements on the used dFBA and experimental setup might solve some of these shortcomings by integrating additional genetic information. This could be used to create a simulation that mimics the situation with a mutation background. Ultimately, the introduction of genetic information with rules that constrain the simulation set even more difficulties to interpret the results. The simulation is required to use highly curated genetic information about growth changing mutations if it should be used for dFBA. Addressing the prior mentioned issues will yield valuable insights into growth, physiology, and metabolism.

In conclusion, our findings support the potential of integrating physiological data into metabolic models enabling the generation of valuable growth-related predictions. This study demonstrated that the physiology of complex multicellular organisms stands in relationship with metabolic processes. We could present that gut growth play a role as a regulatory factor for overall organismic growth, a phenomenon observed in wild-type *D. melanogaster* larvae. The enhanced metabolic model of FlySilico enabled the successful prediction of larval growth over the entire development for various conditions. The incorporation of physiological data from the larval gut enhanced the prediction power of FlySilico, allowing the accurate estimation of developmental critical milestones, including the reach of the critical weight and further the pupation. This enables further in-depth investigations into the mechanisms of critical weight. The extension of this approach to incorporate additional physiological data, potentially involving multiple organs concurrently, holds great promise. These extensions empower the possibility of predicting more favorable growth scenarios.

Ultimately, this approach has the potential to facilitate the translation of these findings to predict even the growth of more complex organisms. Furthermore, this could lead to the possibility to formulate a hypothesis that gut growth and physiology is not only limiting factors for complex organisms like *D. melanogaster* but may also play a crucial role in regulating development and health of extrauterine growing organisms, including humans. We propose that variations in gut physiology, nutrient absorption, and growth in stages, like infants, have a lasting impact on the fitness of individuals and could restrict growth. This hypothesis, along with potentially others, supports future research studies.

## Material and Methods

### Fly strains and rearing

The fly lines that were used in this study are Oregon-R. Flies were maintained at 25 °C with 60-70 % humidity and a 12 h light/dark cycle.

### Chemically defined fly medium holidic diet (HD)

The chemically defined fly medium HD was used based on the instructions of Piper et al. (2014). The parametrization was performed according to Schönborn et al. (2019).

### Complex fly medium low sugar diet (LSD)

The complex fly medium LSD was used based on the complex medium from Backhaus B. et al. (1984) and altered according to Musselman et al. (2011). The parametrization was performed according to Schönborn et al. (2019) and based on the available data of the used ingredients (see Supplementary Table 2). As not all information were unambiguously, we used approximated values.

### Metabolic profiling

All data were measured according to the protocols by Schönborn et al. (2019) and under standard conditions on the growth medium *holidic diet* (Piper et al., 2014; Piper et al., 2017) or low sugar diet.

### Fixation, histological staining, and mounting

The larval guts were dissected in ice-cold PBS and fixed in RNA-fix (10 % 10x PBS, 10 % 0.5 M EGTA pH 8.0, 10 % Paraformaldehyde, dH20 70 % -1:1 dilution before use with dH20) for 40 minutes. The RNA-fix solution was removed, and the guts were washed with PBS (1x) for 5 minutes.

Alexa Fluor® 488 Phalloidin (300 U dissolved in 1.5 ml methanol) was used as the staining agent and diluted 1:500 in PBS before direct usage. 1 ml of the diluted Alexa Fluor® 488 Phalloidin solution were used to stain the guts for 30 minutes. After staining the guts were washed with PBS (1x) for 5 minutes.

The guts were mounted on microscope slides in 30 μl Prolong Gold Antifade reagent.

### Constraint-based modeling

Flux balance analysis (FBA; Orth et al., 2010) was used to perform the underlying growth simulations and metabolic flux analysis.

### Biomass coefficients

We measured the same metabolites in this publication and according to the previously published FlySilico (Schönborn et al., 2019) version. The reason is that the measured metabolites are the most abundant metabolites present in a *Drosophila* larva which should sufficiently represent the growth of the larva. We extended the range where we measured the metabolite contents in comparison to the previously published FlySilico version. Here, we calculated a **<u>w</u>**eighted **<u>m</u>**ean **<u>m</u>**etabolite **<u>c</u>**oefficient (WMMC) for the biomass function. The WMMC should realistically represent a single value of the changing quantity of metabolites to make up the biomass through the larval development. The WMMC is calculated as followed by starting to calculate a polynomial fit *f_x_*(*t*) of the measured data, where *x* represents the metabolite and *t* the timepoint. Next, we used the values of the derivation *f’_x_*(*t*) at the timepoints where we measured our data of the metabolites and normalized them between 0 and 1 as followed:

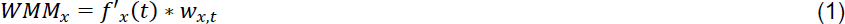

where *WMM_x_* is the **<u>w</u>**eighted **<u>m</u>**ean of **<u>m</u>**etabolite *x* in either *μg* or *nmol* and *w_x,t_* is the measured weight of metabolite *x* at timepoint *t*. With the WMM from Equation 1 we can calculate the WMMC:

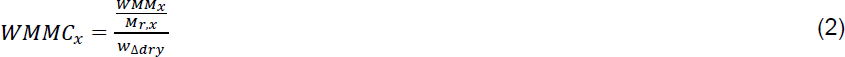

Where *WMMC_x_* represents the weighted mean metabolite coefficient of metabolite *x* in, 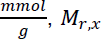 is the molecular weight of metabolite *x* and *w_Δdry_* is the mean weight gain in *g*. The mean weight gain is calculated as followed:

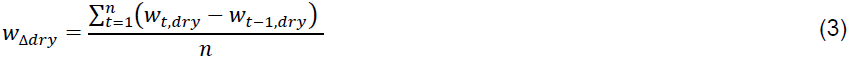

where *w_t,dry_* is the dry weight of the larva at timepoint *t* and *w_t-1,dry_* is the dry weight of the larva at the previous timepoint *t* - 1.

Additionally, we calculated a chitin biomass coefficient based on data present in the literature. The literature regarding the chitin content of larval *Drosophila* was not clear how large it is. Therefore, we used the chitin content of adult flies to approximate the content of larval chitin in combination with our data. Kaya et al. (2016) determined the content of chitin in adult fly is 7.85 %.

The chitin biomass coefficient was calculated with the following equation:

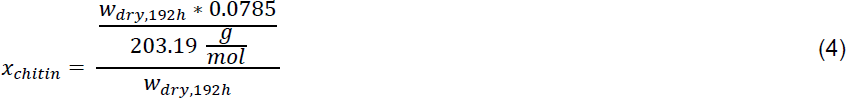

where *x_chitin_* is the biomass coefficient of chitin represented in the biomass function, *w_dry,192h_* is the dry weight in gram dry weight (*gDW*) of a larva at 192 hours AEL on HD, 0.0785 represents the chitin content from literature (Kaya et al., 2016) and 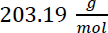 is the molar mass of chitin.

The resulting chitin biomass coefficient is 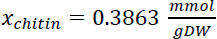.

### Growth model equations

The larval growth of *Drosophila* is described by literature as an exponential growth. The growth in our model according to the literature modeled with a basic exponential growth function:

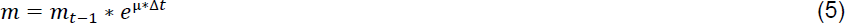

where *m* describe the biomass of the larval at the timepoint *t* in *m_gDW_, m_t_*_-1_ describes the biomass of the larval at the previous timepoint *t* − 1, µ describes the growth rate derived from the FBA solution and Δ*t* describes the timestep size for which the calculation is.

The uptake rates *v_x,t0_* of each most abundant metabolite *x* in the growth medium HD at the initial timepoint *t_0_* where calculated based on the new experimental data according to Schönborn et al. (2019).

The uptake rate at each timepoint in the dynamic FBA (dFBA) approach is calculated as followed:

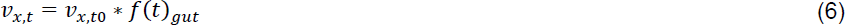

where *v_x,t_* describes the uptake rate of metabolite *x* at the timepoint *t* in 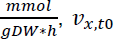 describes the uptake rate of metabolite *x* at the initial timepoint *t*0 and *f*(*t*)*_gut_* describes the dimensionless gut resorption penalty function.

The gut resorption penalty function is defined as following sigmoidal function:

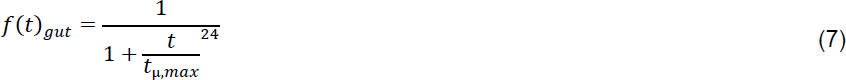

where *t* describes the current timepoint in the dFBA simulation in hours and *t*_µ,*max*_ describes the timepoint where the growth of the larvae should be at maximum in hours. Therefore, *t*_µ,*max*_ is defined as followed:

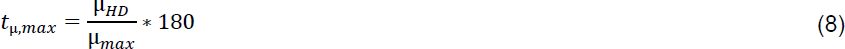

where µ_*HD*_ describes the growth rate for larvae on HD growth medium in *h*^−1^, µ_*max*_ describes the maximum growth rate of the current simulation time range and 180 hours describes marks the timepoint where the larval gut of larva grown on HD medium enters the plateau-like state (Fig. 3 D-F). Furthermore, this timepoint lies within the range when the larvae attain the critical weight threshold while fed on HD medium.

## Acknowledgements

We would like to thank all members of the laboratory for their helpful comments and support, and Oliver Ebenhöh for his discussions during the initial phase of the project.

## Contributions

J.S. and M.B. designed the study. J.S. established the metabolic network and performed the modeling experiments. J.S., A. D., I. A., and S. W. performed the *Drosophila* experiments and wet lab procedures. J.S. and

M.B. analyzed the data and prepared graphs. J.S. and M.B. wrote the manuscript.

---

**Supplementary Figure 1:**
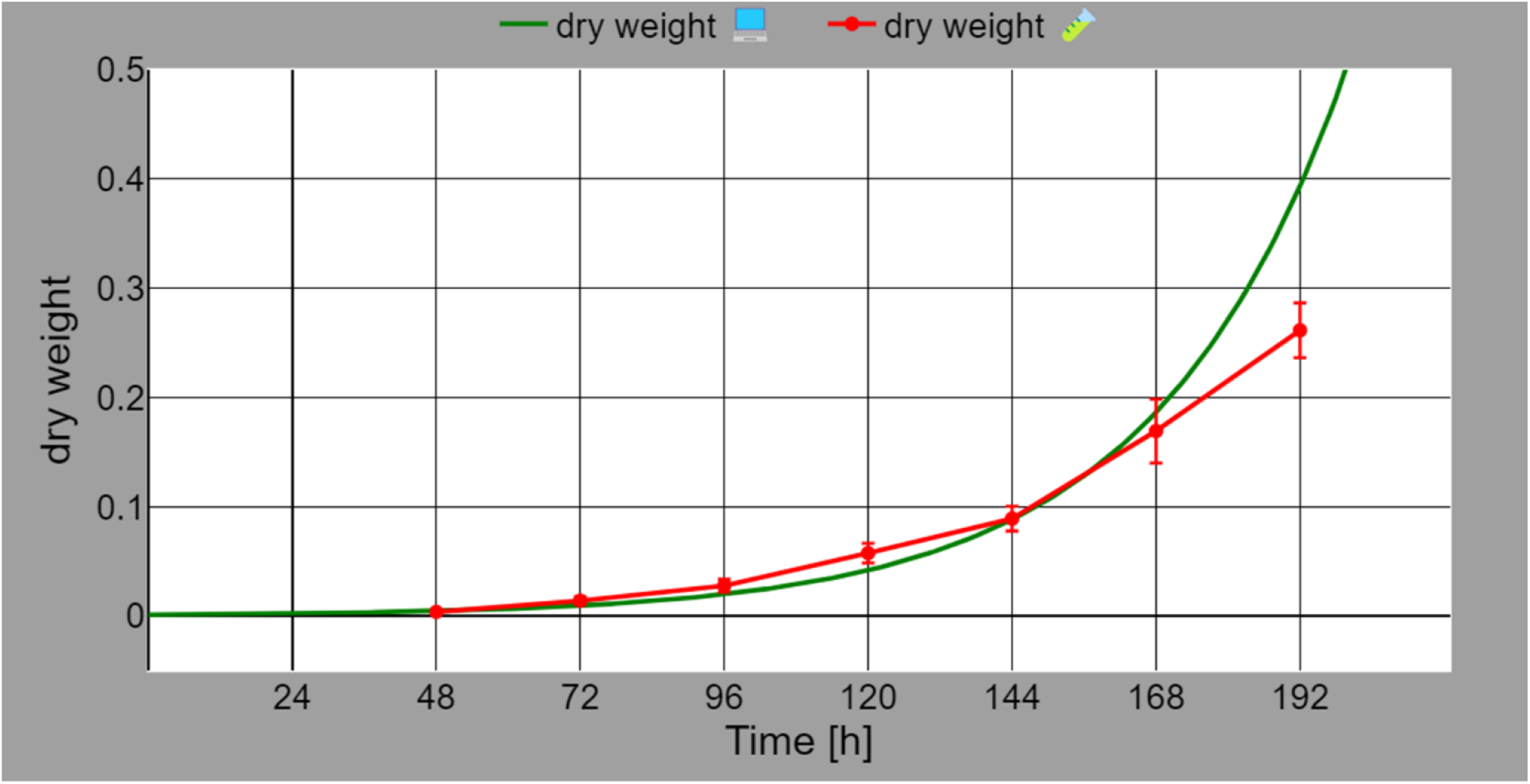
Comparison of *in vitro* and *in silico* larval growth on chemically defined growth medium. We simulated the *in silico* growth of a larva (green; dry weight) on HD by using the dFBA approach (see method section) with the addition of the penalty function. The growth of the larva resembles a exponential growth curve with unlimited growth. The growth of larvae reared *in vitro* (red; dry weight) on HD, starts at 48 hours till 192 hours AEL. The growth *in vitro* shows a small variance in comparison to the *in silico* growth till it reaches 168 hours AEL. On average the pupation started around the 192 hours mark.

**Supplementary Figure 2:**
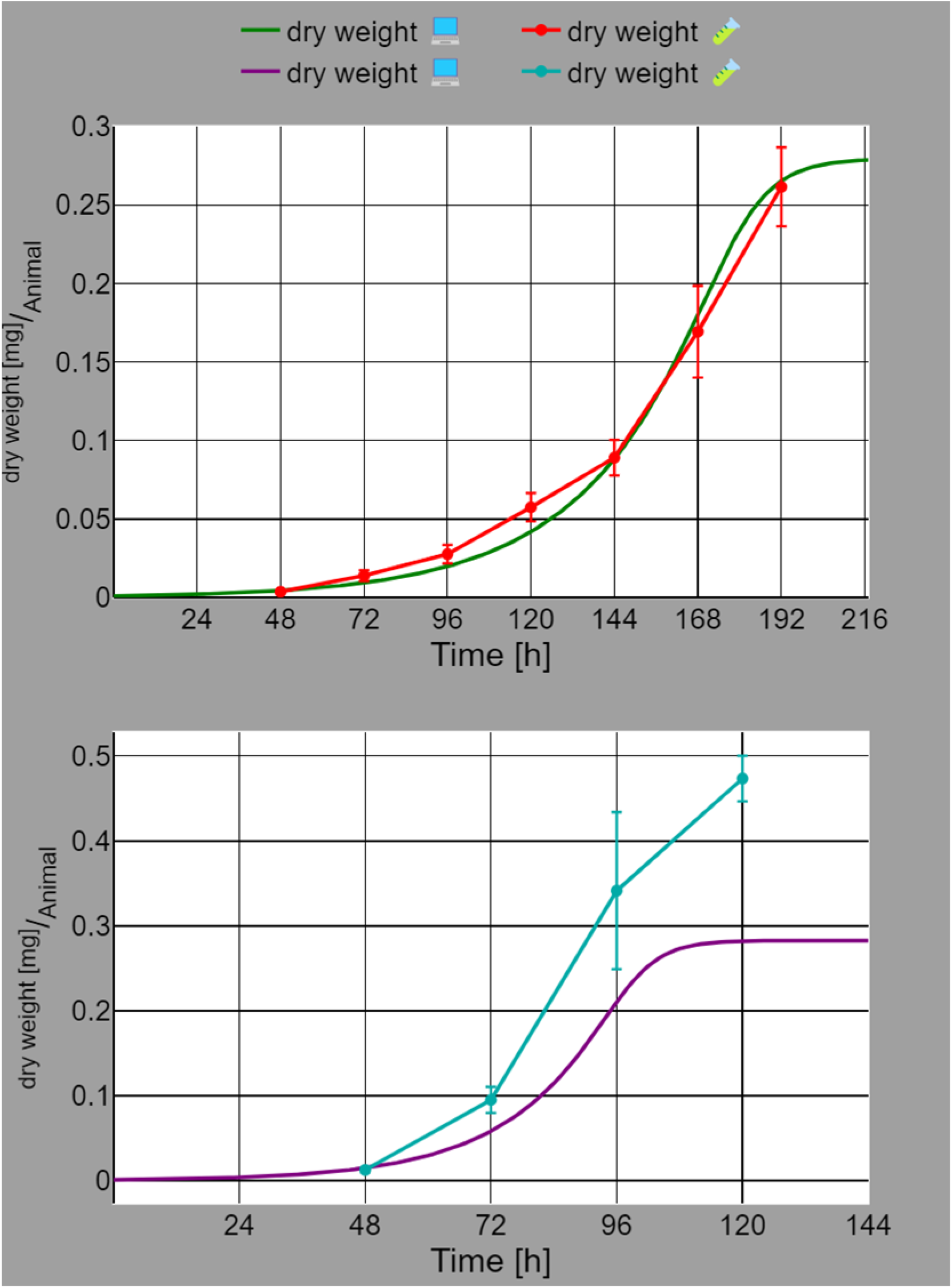
Comparison of *in vitro* and *in silico* larval growth on chemically defined growth medium and complex medium. (A) We simulated the *in silico* growth of a larva (green; dry weight) on HD by using the dFBA approach (see method section). The growth of the larva resembles a sigmoidal growth curve till it reaches a plateau starting around 192 hours. The growth of larvae reared *in vitro* (red; dry weight) on HD, starts at 48 hours till 192 hours AEL. The growth *in vitro* shows a small variance in comparison to the *in silico* growth based on the absolute dry weight values per animal. On average the pupation started around the 192 hours mark. (B) We simulated the *in silico* growth of a larva (purple; dry weight) on HD by using the dFBA approach (see method section). The growth of the larva follows a sigmoidal growth curve till it reaches a plateau starting around 120 hours. Larval growth observed *in vitro* (dark cyan; dry weight) on HD medium, starts from 48 hours till 120 hours AEL. The *in vitro* growth shows a greater variance when compared to the *in silico* growth data based on the absolute dry weight values per animal. While the *in silico* results do not match the absolute values observed *in vitro*, they do show a similar growth trajectory as they are aligning with the correct timing for the conclusion of growth.

**Supplementary Figure 3:**
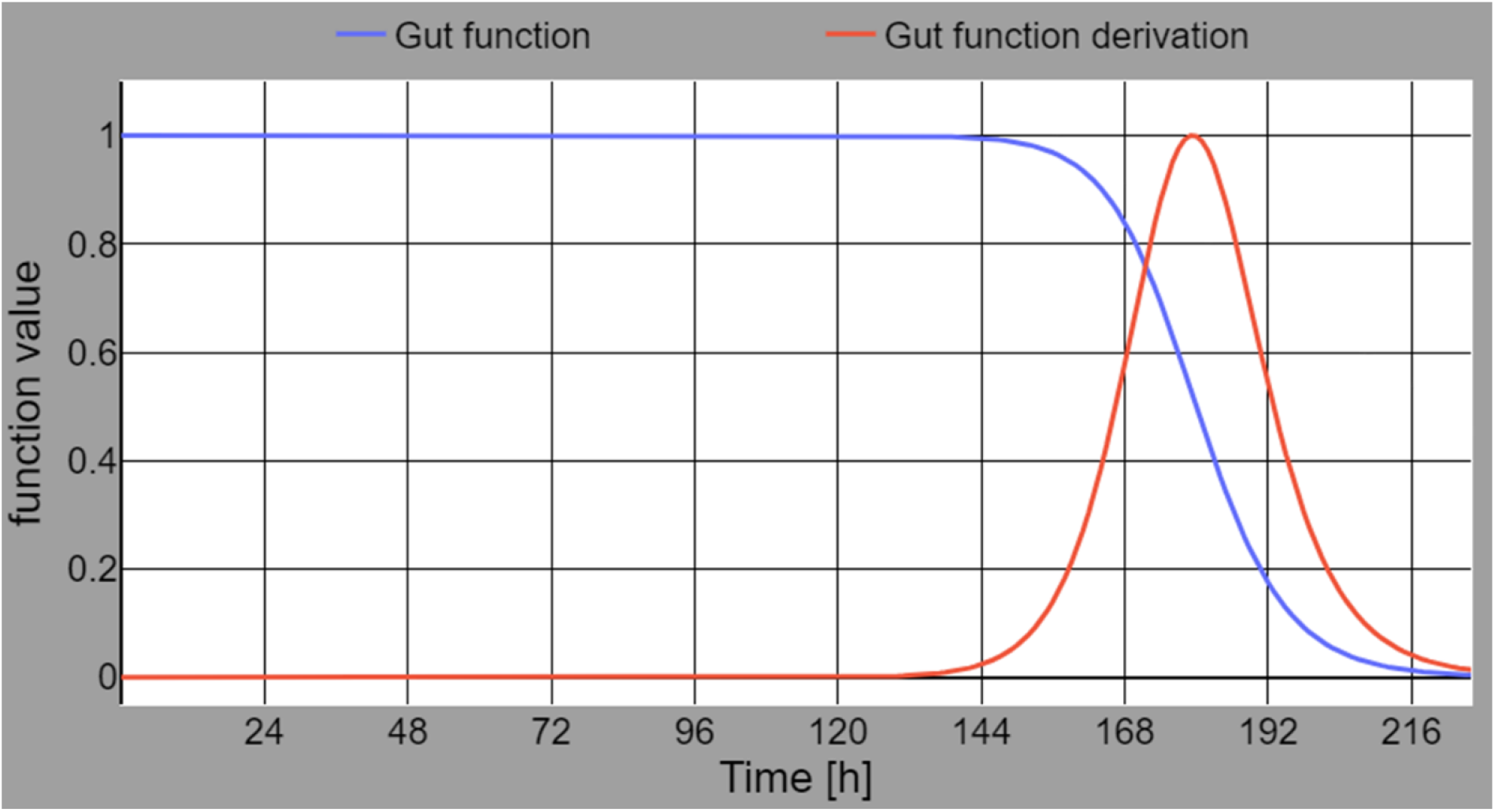
Penalty function of the *in silico* simulation based on HD. The penalty function or “gut function” (blue) was formulated as described in the method section. From 0 hours till 128 hours the function value is 1. The gradient starts to change from 128 hours till 218 hours till the function value of nearly 0. The turning point of the function is at 180 hours with a function value of 0.5. The “gut function derivation” (red) is the derivation of the gut function (blue). It shows a function value change in the phase of the gut function where the gradient starts to change. The maximum function value is 1 at 180 hours which is the turning point of the gut function. For better representation the “gut function derivation” values are normalized absolute values.

**Supplementary Figure 4:**
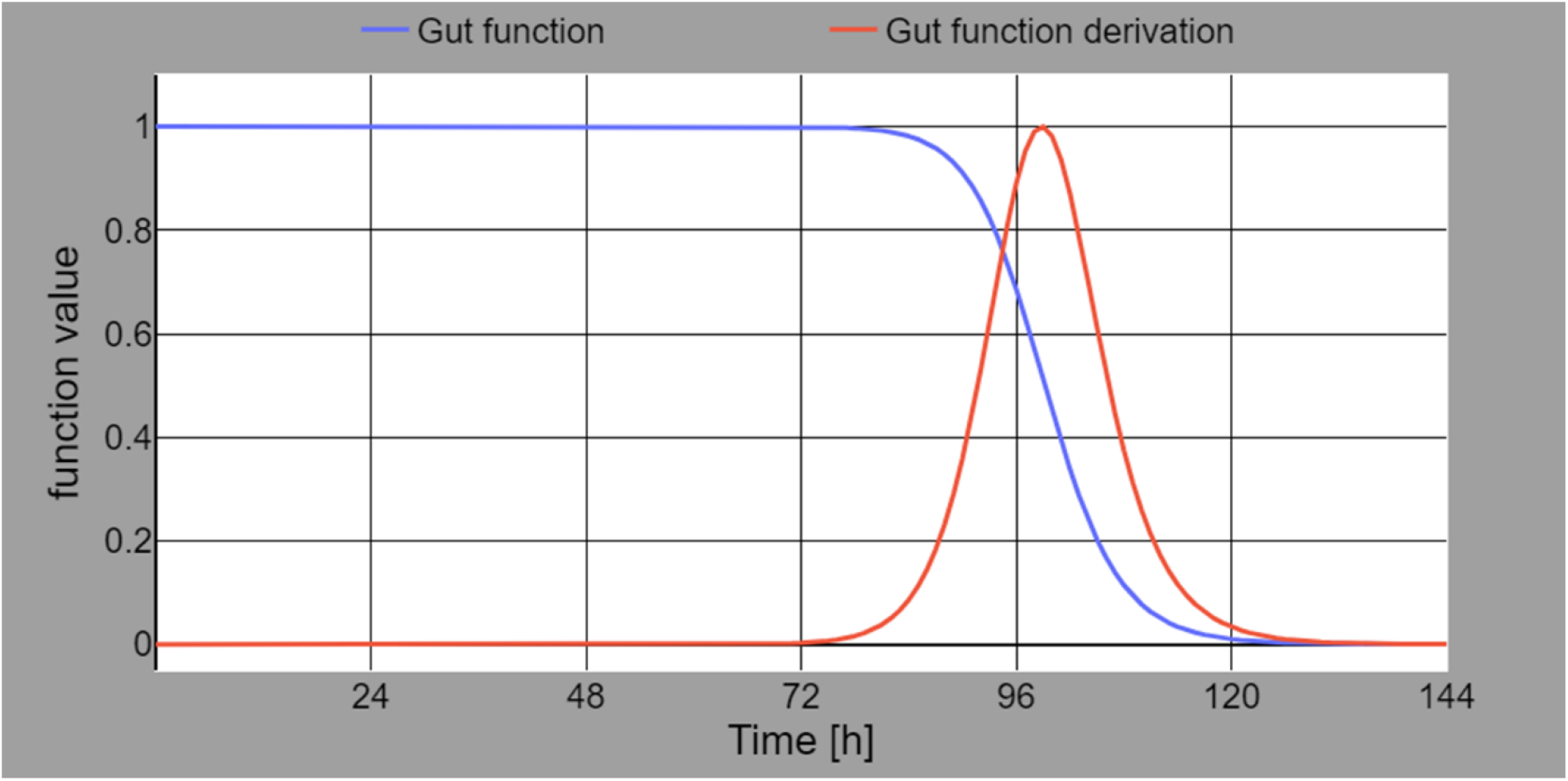
Penalty function of the *in silico* simulation based on LSD. The penalty function or “gut function” (blue) was formulated as described in the method section. From 0 hours till 76 hours the function value is 1. The gradient starts to change from 76 hours till 128 hours till the function value of nearly 0. The turning point of the function is at 100 hours with a function value of 0.5. The “gut function derivation” (red) is the derivation of the gut function (blue). It shows a function value change in the phase of the gut function where the gradient starts to change. The maximum function value is 1 at 100 hours which is the turning point of the gut function. For better representation the “gut function derivation” values are normalized absolute values.

